# Report on the spatial spread of defective interfering particles and its role in suppressing infectious particles

**DOI:** 10.1101/558932

**Authors:** Qasim Ali, Ruian Ke

## Abstract

Defective interfering particles (DIPs) are categorized as non-infectious viruses with large deletions in their genomic material. A cell infected by a DIP require co-infection by a wild-type virus to complete its replicative lifecycle. There is an increasing interest in developing DIP based therapies in the form of molecular parasites that steal genetic resources of infectious particles. This parasitic behavior is enhanced by constructing engineering designs of DIPs to optimize their role in suppressing the virus infection within-host. Recent experimental studies characterize viral infection as a spatial process and emphasize on its spread rate and the area populated by the infectious particles (IPs). We developed a spatio-temporal model in the framework of reaction-diffusion equations to depict the functional organization of virus particles distributed over a tissue surface. Our model investigates the scenarios and figures out the aspects that can play a vital role to suppress the infection within-host. We studied the impact of initial dose of DIPs, the efficiency of DIP production and the role of cell maturation. Our results show that an engineered DIP can substantially decrease the concentration of IPs. We assert that the decrease in the rate of spatial spread of IPs requires non-deterministic settings.

## Methods

### A spatial model for virus infection including defective interfering particles

We construct a spatial model using partial differential equations (PDEs). We consider six populations of cells and two populations of virus. These populations of cells include target cells *T*, three eclipse phases for non-productive infected cells (*D, E* and *E*_D_), two classes of infective cells (*I* and *I*_D_) and two virus populations (*V* and *V*_D_). The system of PDEs is as follows:

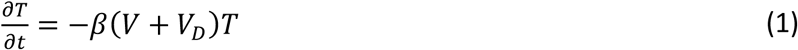

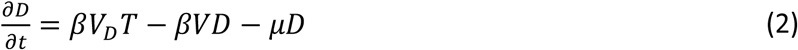

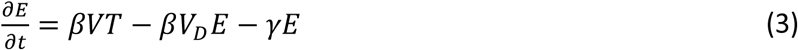

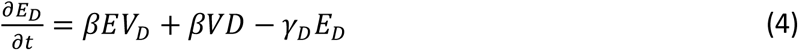

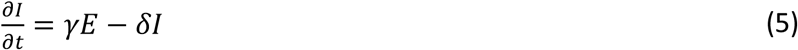

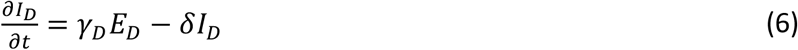

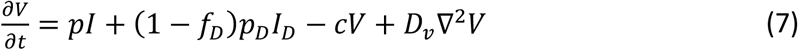

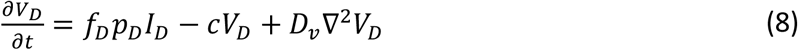

with initial and boundary conditions of the system as:

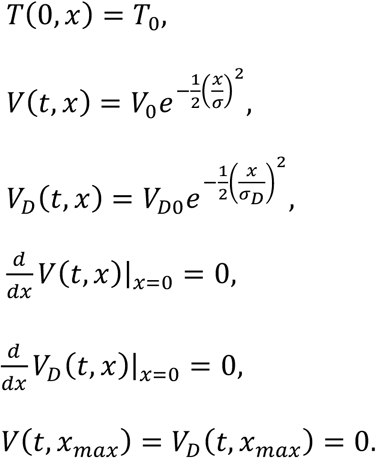

and

In this model, virus particles are initially spread on the spatial domain *x* ≥ 0 where target cells are homogeneously present throughout the space. Reaction-diffusion modeling framework is used to define the spread of virions and cells while the virus particles are allowed to diffuse. Population densities of the virus particles are represented by *V*(*x,t*) for infectious particles (IPs) and *V*_D_(*x,t*) for defective interfering particles (DIPs).

Viruses infect target cells *T* (cells) at infection rate constant *β* (TCID_50_/ml) ^−1^ day^−1^. Infections caused by DIPs to the target cells *T* results in the increase of defective-infected cells in the class *D*. These cells are unable to produce infectious particles; they require co-infection by IPs in order to produce the virus particles *V* and *V*_D_. However, this is only possible during the first few hours of infection, i.e. the incubation period, where there is no virus production. The DIP infected cells become matured at the rate *μ* day^−1^ and no further infections are possible. The target cells infected by IPs are represented by an eclipse class (*E*) where co-infection by DIPs is possible during their incubation period. The co-infections to the cells in *E* class by the DIPs forms a new eclipse class (*E*_D_) at the rate *β* virion^−1^ day^−1^.

The cell populations in the classes *D, E* and *E*_D_, are non-productive, however maturation of *E* and *E*_D_ classes lead to the productive compartments. Cells in *E* and *E*_D_ matures at rate *γ* and *γ*_D_ (day^-1^) to become productively infective cells of classes *I* and *I*_D_ respectively. The infective class *I*_D_ will produce DIPs and IPs whereas infective class *I* can only produce IPs. The cells present in the infective classes die at rate *δ* (day^-1^). Virions production rate from cells in *I* and *I*_D_ classes are denoted by the constant rates *p* and *p*_D_ (TCID_50_/ml) cell^-1^ day^-1^ respectively. We define *f*_1_ as a fraction of virions produced by *I*_D_ class that are DIPs. In this way, rest of the (1 – *f*_1_) virions produced by *I*_D_ class are considered IPs. The viruses have the ability to diffuse in the mucosa present around the cells with the rate of diffusivity *D*_v_ whereas the viruses get cleared from the system at a constant rate *c* (day^-1^).

This study aims to understand the qualitative impact and general features of DIPs on the spatial viral spread, instead of focusing on specific parameter values, however, the baseline parameter values are defined to represent biologically realistic dynamics (see Table 1). The initial population of target cells is homogeneously distributed over the space: *T*(0, *x*) = *T*_0_ (*x*) = 6.6 x 10^5^ cells/cm^2^. The initial distribution of IPs and DIPs has same baseline maximum initial values *V*_F0_ = *V*_D0_ = 1 and same variance for initial spread of virions *σ*_F_ = *σ*_D_ = 0.1.

**Table 1:**
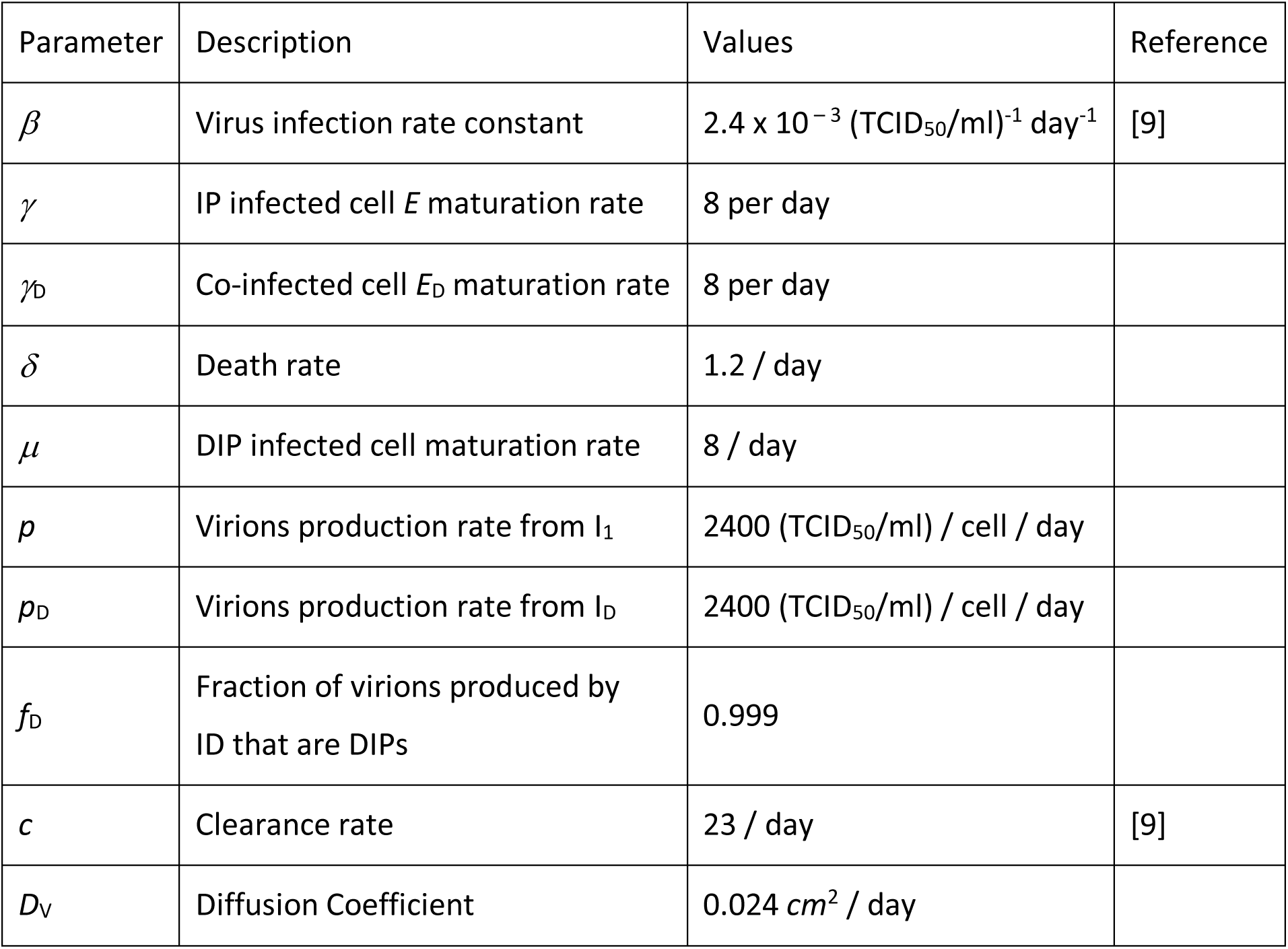
Parameters and their values used in this study

Initial phase of infection is hard to estimate in-vivo; however, in-vitro study provides some insight to the spatial aspects of infection (Gog et al., 2015). In regards to spatio-temporal models for within host viral dynamics during first day of infection, there is very lite information known about infection rate. Therefore, the parameter for the infection rate constant *β* is chosen so that virus spread in the absence of DIP. Viruses are able to diffuse on the surface of epithelial layers during the respiratory tract infection (van Riel et al., 2007).

A study provides estimate for the diffusivity of virions in spherical shape with diameter 10 to 200 nm in the interval [0.0016, 0.026] cm^2^/day in plankton (Murray and Jackson, 1992). Several other studies also considered the virions diffusivity in the consistent range (Chang et al., 2008; Hu et al., 2011; Schmid et al., 2015). Diffusivity of virions through the mucus depends upon the spacing between mucin fibers, their diameter and the size of virions. Typical size of influenza virus (between 80 and 120 nm) can diffuse almost unhinderedly through such fibers due to large mesh spacing between the mucin fibers (Olmsted et al., 2001). Therefore, we set the diffusivity constant within the given range as 0.024 cm^2^/day to make sure virus particles can diffuse easily.

Parameters that are associated to single cell dynamics, e.g. maturation rates of infection *γ, γ*_D_ and *μ*, and death rate of infected cell *δ*, are extensively studied in the literature. Since cells are not diffusing in the system and homogeneously distributed over the whole spatial domain, it is easy to follow these rates from the literature (Baccam et al., 2006; Beauchemin and Handel, 2011; Canini and Carrat, 2011; Pawelek et al., 2012). The virions clearance rate *c* is another extensively studied parameter and widely accepted same as defined with 95% confidence interval for the acute phases of viral infections (Baccam et al., 2006; Beauchemin and Handel, 2011; Cho et al., 2014; Perelson et al., 1996; Smith and Perelson, 2011). A recent study shows that virions clearance rate is highly dependent on the density of infected cells (Smith et al., 2018). However, during the first day of infection, the virions either quickly infect cells or get cleared from the host. This allows us to calculate virus clearance rate by counting number of dead cells during early period of infection.

Virions production rate per cell (*p* and *p*_D_) and fraction of DIPs produced by matured DIP infected cell (*I*_D_) are the parameters of analyses for this study. These parameters are significant to design the drug as their variability can directly affect within host virus infection.

## Results

We constructed a spatial model in the framework of a reaction-diffusion system of partial differential equations to describe the impact of DIPs on the virus spread. Through the model we explore interference of DIPs in the spread of infectious particles and their role in suppressing infection in spatio-temporal domain. A schematic for the mathematical model is presented in the Figure 1.

**Figure 1:**
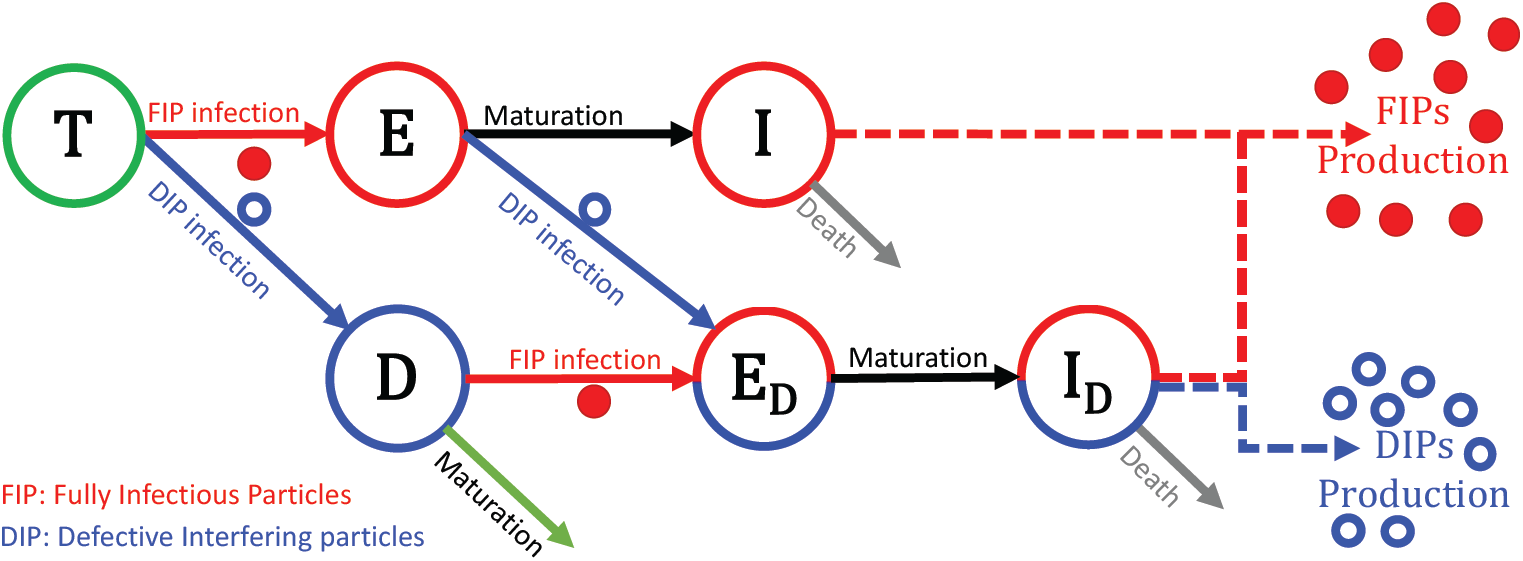
A schematic representation of the diffusion model. Non-infectious classes are target cells *T*, eclipse phase for cells infected by IPs *E*, eclipse phase for cells infected by DIPs only *D* and eclipse phase for cells infected by DIPs and IPs *E*_D_. Infectious classes are IPs producing cells *I* and IPs and DIPs producing cells *I*_D_.

### Possible scenarios for the survival of IPs and DIPs

Following an initial period of transient growth dynamics during early infection, the spread of infection takes the form of a constant-speed traveling wave solution. The transient phase is dependent on the initial conditions of virus populations and the production of virus particles by infective cells. A necessary condition for the production of DIPs from the infective class *I*_D_ is the occurrence of co-infection with both DIPs and IPs, whereas production of IPs does not require any co-infection and it alone can maintain the infection process and establish a traveling wave front. Because of this, the establishment of a sustainable level of DIP production during the initial transient phase of infection is dependent on the ratio of DIP production from co-infected cells. When the ratio of DIPs to IPs produced is small, insufficient initial production of DIPs will not yield a sufficient amount of coinfection for DIPs to attain sustained spatial spread, see Figure 2 panel (a). However, for relatively high values of *f*_D_, DIPs survival is possible. With the co-survival of virus particles and increase in the production of DIPs from co-infected cells, IPs decrease. This decrease is observed in the peak but not in the speed of the spatial spread of infectious particles *V*_F_ (see Figure 2 panel (a) and (b)).

**Figure 2:**
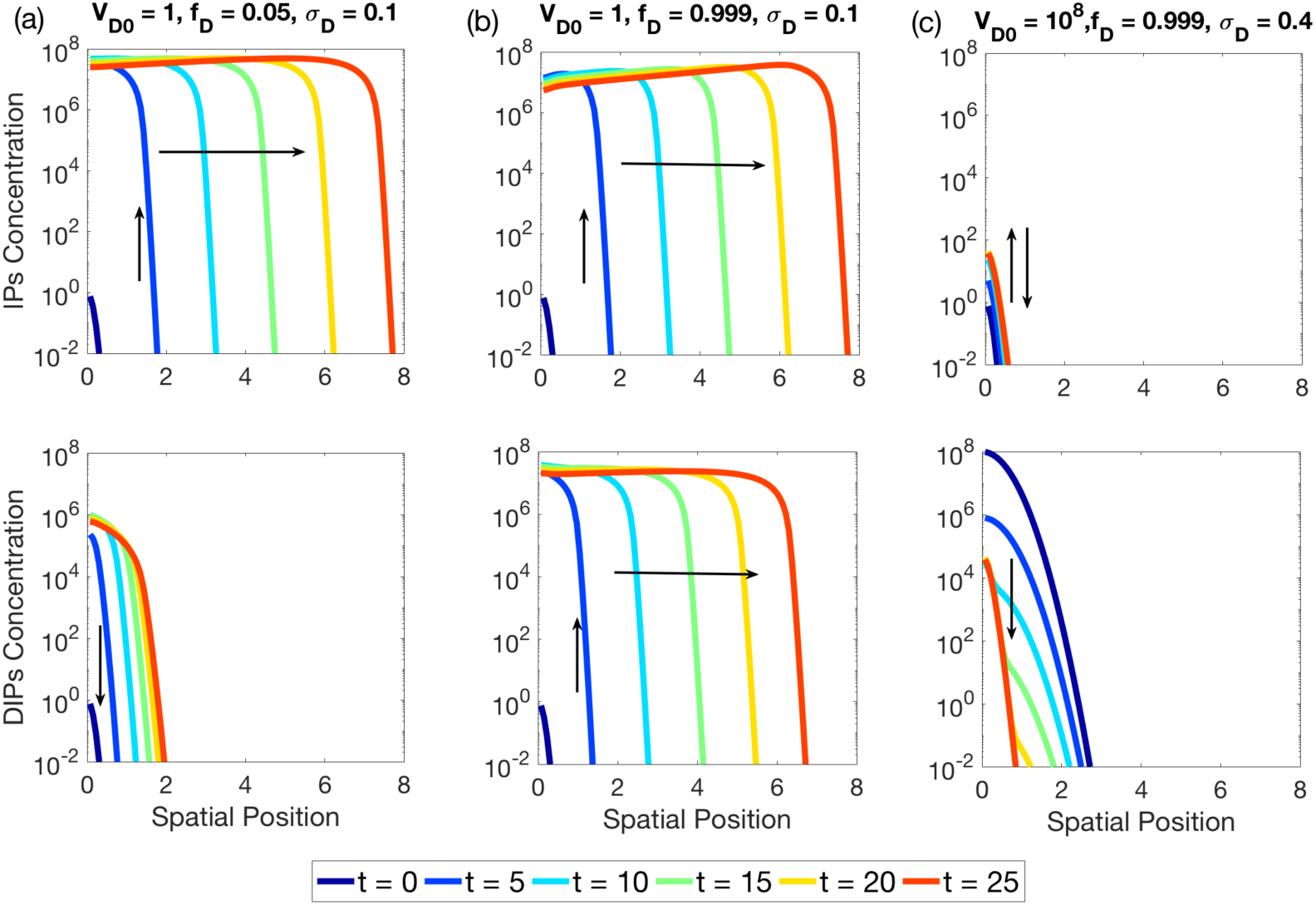
Each curve represents concentration of IPs and DIPs spread over the spatial domain at specific time. Three panels are drawn; the plots in the top row in each panel show the spread of fully infectious particles (IPs) whereas the plots in the lower row represents the spread of defective interfering particles (DIPs). **(a)** Time is varied from 0 to 40 hrs. At time *t* = 0, maximum initial concentration and initial variance of DIPs and IPs are chosen same (*V*_D0_ = *V*_F0_ and *σ*_D_ = *σ*_F_) while fraction of DIPs produced from the cells in *I*_D_ class are near their lowest value (*f*_D_ = 0.05), **(b)** Time is varied from 0 to 20 hours. At time *t* = 0, *V*_D0_ = *V*_F0_ and *σ*_D_ = *σ*_F_ and *f*_D_ = 0.999, **(c)** Time is varied from 0 to 80 hours. At time *t* = 0, *V*_D0_ = 10^8^ *V*_F0_ and *σ*_D_ = 4*σ*_F_ and *f*_D_ = 0.999.

During the early period of infection, we observe that DIPs need to attain sufficiently high peak value in a limited amount of time in order to propagate along IPs. This is necessary in a time period when IPs have not attained a constant speed at the travelling front. In the case of choosing a high initial peak value (*V*_D0_ ≫ *V*_F0_), and characteristic width (*σ*_D_ ≫ *σ*) of the initial distribution of DIPs so that a substantial amount of local target cells is infected by DIPs, the infection process can be halted. Consequently, due to the requirement of co-infection by the DIPs infected class *D* to become infective class *I*_D_, the DIPs spread will also be reversed, see Figure 2(c).

In Figure 2, the IPs are reduced by increasing the fraction parameter *f*_D_ and the initial condition for the DIPs. We incorporate these observations to our subsequent results in order to investigate their impact on reducing *V* by decreasing the travelling wave speed of the front as well as its peak value.

### DIPs role in reducing infectious particles

During the first few hours of infection, when the DIP concentration is low, increase in IPs is only due to its production from infective class *I*. Therefore, the impact of *f*_D_ is initially minimal. During this time, IPs reach their peak value and form a travelling wave front as shown in Figure 3 (a), (b) and (c). We observed in Figure 2 that when the initial conditions are same for both virus particles, the IPs lead the travelling wave front. Therefore, the speed of the travelling front remains unaffected by the variation of *f*_D_ since the speed of propagation is determined by the infection dynamics at the leading edge of the front, as shown in Figure 3((g), (h) and (i)). During this initial period when the system is exhibiting transient dynamics, the peak value of IPs remains stationary as shown in Figure 3(d), (e) and (f). Once productive infection by IPs is established, there are sufficiently many IPs for co-infection to occur and the level of DIPs will begin to increase. As the DIPs come up, the peak value of IPs goes down so that the peak position shifts to the right. Thus, in case of no DIPs, the peak value stays at the initial position a for longer period of time. DIPs are active only locally to where they are produced and are unable to infect target cells which lead the front of IPs, so the front of DIPs will lag behind the front of IPs. One can consider the pointwise dynamics (at a specific spatial position *x*) of the reaction-diffusion system to mimic the time-dependent dynamics of the reaction terms alone.

**Figure 3:**
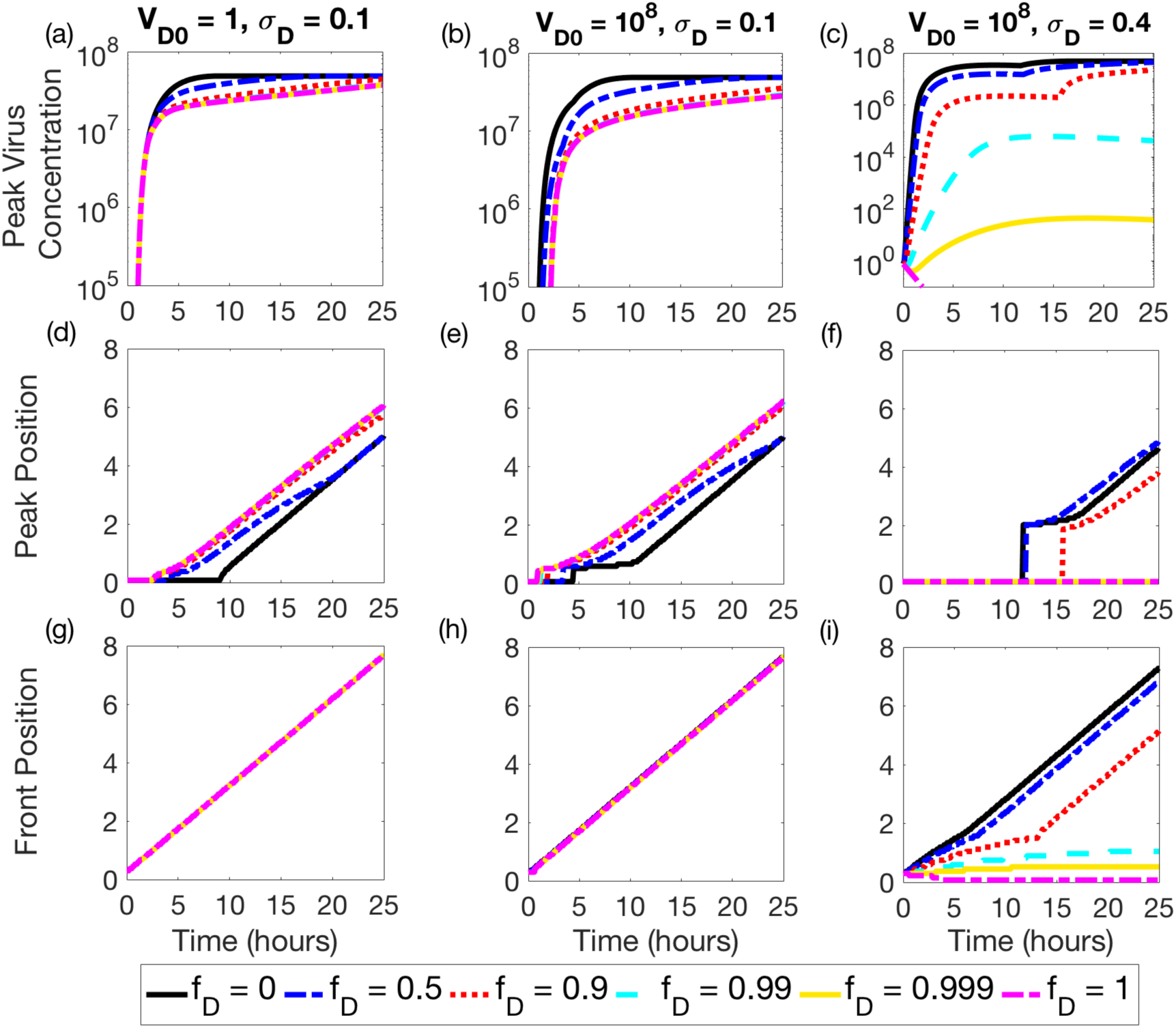
Peak value of infectious particles, peak position and front position of the travelling wave during 20-hour simulation time. Figures are distributed in three columns based on the parameters involved in the Initial condition for DIPs. Parameters in the subplots in column 1 are (a), (d) and (g): maximum initial concentration and initial variance of DIPs and IPs are chosen same (*V*_D0_ = *V*_F0_ and *σ*_D_ = *σ*_F_). Parameter in the middle column has subplots (b), (e) and (h) such that *V*_D0_ = 10^8^ and *σ*_D_ = *σ*_F_. For subplots in column 3, i.e. (c), (f) and (i), the values are *V*_D0_ = 10^8^ and *σ*_D_ = 4*σ*_F_. Each subplot contains six curves at distinct values of *f*_D_. These values are *f*_D_ = 0 (black solid line), 0.5 (blue dash-dotted line), 0.9 (red dotted line), 0.99 (cyan dashed line), 0.999 (yellow solid line) and 1 (maroon dash-dotted line) are chosen for each plot.

The spatial virus dynamics are different if the initial conditions are varied. For high parameter values of DIPs initial distribution, DIPs infect more target cells at the early spatial position. This results in the increase of cells in the eclipse class *E*_D_ rather than *E*. Thus, for high values of *f*_D_, when more DIPs are produced from the cells in *I*_D_ class, the peak value of IPs either remains at early spatial position and diminishes with time or emerges late in time from the wave front, see Figure 3(last column). This looks like a discontinuity as shown in Figure 3(f).

The speed of the travelling front does not show any significant effect by the variation of *f*_D_ unless the peak value as well as the characteristic width of the initial distribution is high, see Figure 3(bottom row). As the cells in the infective class *I*_D_ arise, the role of *f*_D_ becomes prominent. Consequently, the IP production is shared between the two infective classes and the impact of *f*_D_ becomes visible in the form of IPs reduction, see Figure 3 (a), (b) and (c).

The magnitude and characteristic width (*V*_D0_ and *σ*_D_) and of the initial spatial distribution of DIPs plays a critical role in shaping the course of the transient growth phase and determining whether or not a traveling wave is established. However, increasing either of the two parameters independently does not yield any observable effect on travelling front of *V* for any *f*_D_. For instance, maximum initial concentration of DIPs sufficiently higher than IPs is unable to show any prominent effect on the spatial position of infectious particles as shown in the Figure 3(h). This shows that a successful establishment of DIPs requires a large and widely distributed initial DIP population (see Figure 3(c), (h) and (i)). In this way, for high values of *f*_D_, the total IPs and the speed at the travelling front substantially decreases during the whole simulation time.

### Infectious particles over the parameter space spanned by *f*_D_ and *V*_D0_

The transient behavior of virus particles is essential to understand the initial dynamics of infection. However, the net result lies at the end of transient phase when the virus particles establish a travelling wave front. In Figure 2, we observed that DIPs eradicate IPs when their initial characteristic width is higher than IPs (*σ*_D_ ≫ *σ*_F_) if the magnitude of DIPs initial distribution (*V*_D0_) and the fraction of DIPs production (*f*_D_) are sufficiently high. Low DIP production from the *I*_D_ class (*f*_D_ << 1) are unable to affect the IPs, see Figure 4(a) and (b). This is observed even when the initial distribution of DIPs is large and spread out (*V*_D0_ ≫ *V*_F0_ and *σ*_D_ ≫ *σ*_F_). However, DIPs play a good role at the infected region after the early transient behavior where the infected cells in *E* class are co-infected by DIPs. Consequently, the infective cells in *I*_D_ increases and IPs concentration drops down. This shifts the peak position of the IPs near the wave front.

**Figure 4:**
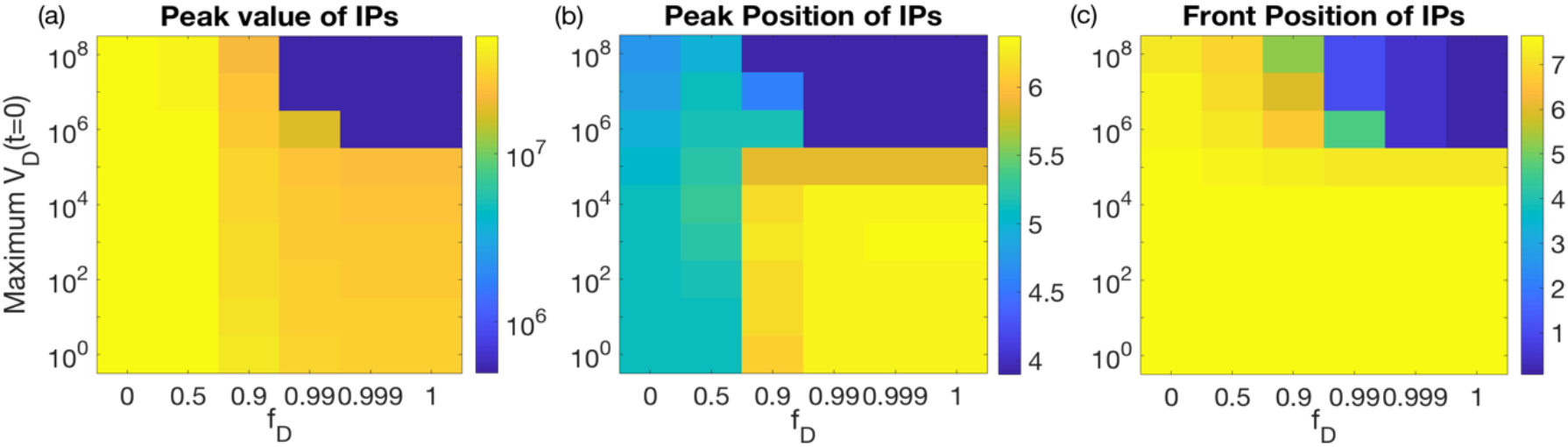
Heatmap for the parameter space *V*_D0_ and *f*_D_ at the final time *t* = 25 hours and *σ*_D_ = 0.4. The spectrum of colors in the color-bar represents the peak concentration of IPs in subplot (a), peak position of IPs in subplot (b) and spatial front position of IPs in subplot (c). The parameter values for fraction of DIPs produced by ID class (*f*_D_) are 0, 0.5, 0.9, 0.99, 0.999 and 1 on the horizontal axis and the values for magnitude of initial DIPs (*V*_D0_) are 10^0^, 10^1^, 10^2^, 10^3^, 10^4^, 10^5^, 10^6^, 10^7^, 10^8^ are on vertical axis. The dark blue region on the top right of each subplot is below the reach of heatmap and therefore matches with the color that is representing the least value in the color map.

High parameter values for the initial distribution reduce the spatial spread of infectious particles in the beginning of the infection because DIPs are able to infect most of the target cells locally. In that case, the production of IPs is driven primarily by cells. In case of high values of *f*_D_, there is insufficient IP production from *to* form a wave front as shown in Figure 4(c). However, for low values of *f*_D_, IPs spatial spread is less affected when the initial distribution of DIPs is high. As the initial distribution of DIPs decreases, IPs are less dependent on cells in *I*_D_ and are able to form their wave front more quickly.

### Production increase from *I*_D_ reduces infectious particles

The fraction of DIPs produced by *I*_D_ class (*f*_D_) played a major role in reducing infectious particles. It is therefore interesting to investigate its efficacy against the number of virus particles produced by the cells in *I*_D_ class, i.e. *p*_D_/*p* = *n-fold increase*. In the above-mentioned results, the production of virions from the cells in *I* and *I*_D_ classes are same (*n* = 1). When the *n-fold* increase is high (*n* > 1), IPs production from *I*_D_ class also increases as shown in Figure 5(a). However, this increase depends on the fraction of DIPs produced by *I*_D_ class. When the fraction of DIPs production is at low, there is a decrease in the number of DIPs infective cells (*I*_D_) and therefore IPs production is solely driven by *I* class. For high fraction of DIPs production, majority of the cells become DIP infective cells.

**Figure 5:**
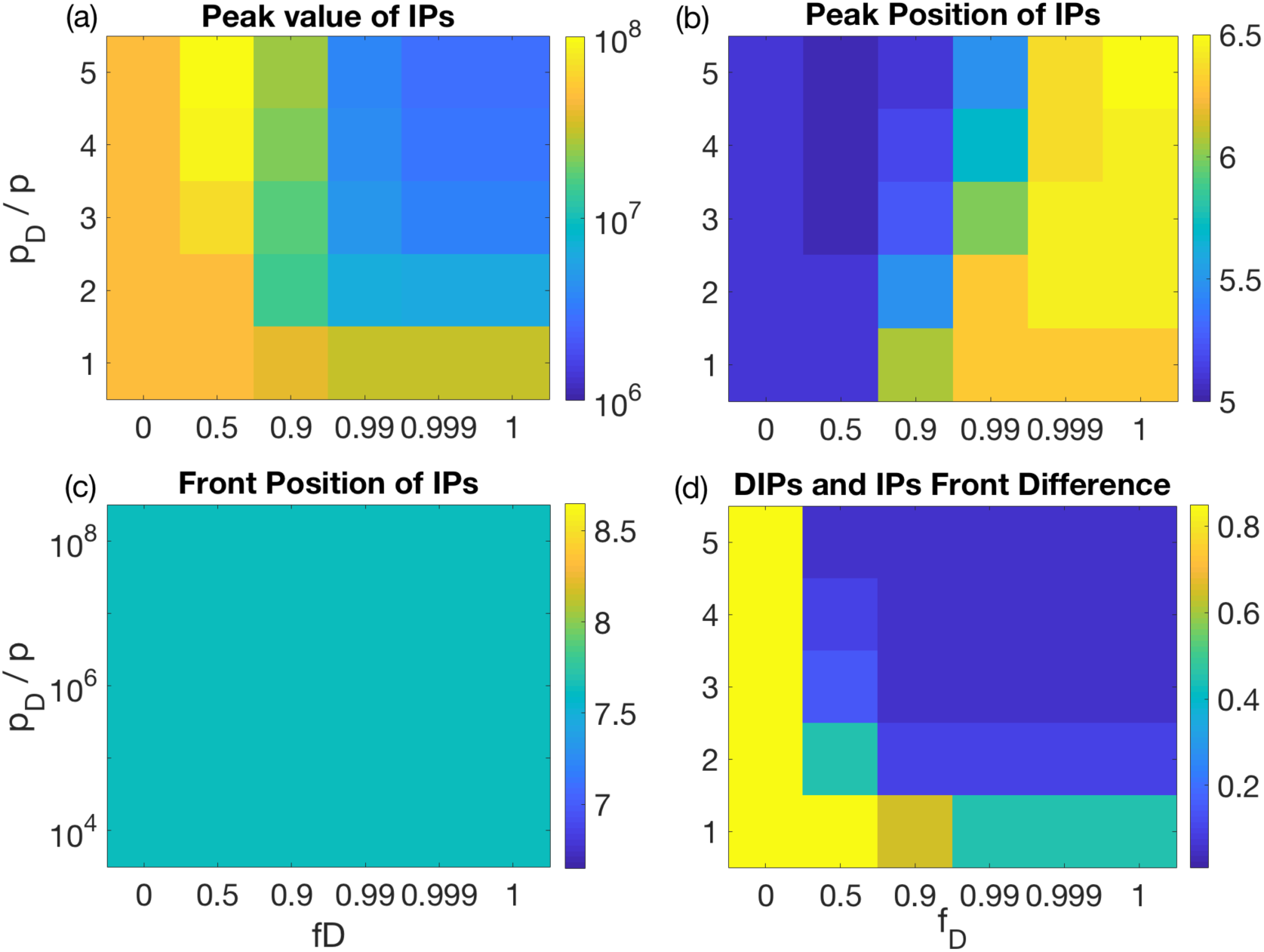
Heatmap for the parameter space, fraction increase in production of virus particles from the cells in *I*_D_ class (n = *p*_D_/p) and fraction increase in the production of DIPs from the cells in *I*_D_ class (*f*_D_), at the final time *t* = 25 hours and *σ*_D_ = 0.4. The color-bars represent the spectrum of colors present in the figures for each combination of parameter values. Subplot (a) describes the parameter space for the peak value of IPs, (b) describes the spatial position of IPs at the peak value and (c) describes the spatial position of IPs at the wave-front. The parameter *f*_D_ values are 0, 0.5, 0.9, 0.99, 0.999 and 1 on the horizontal axis whereas the values for *p*_D_ are 1, 2, 3, 4 and 5 are on vertical axis.

The IPs concentration decreases from the tail end of the travelling wave while its peak shifts near the front position of the infection, see Figure 5(b). We observed that the speed of the travelling front for IPs remain same during the infection period whereas the travelling wave speed of the DIPs increases with the increase in the production of virus particles from *I*_D_ class as shown in Figure 5(c) and (d). The DIPs production increases either due to the increase in the fraction of DIPs production or due to the fold-increase in the virus particles production. The decrease in the difference between the travelling fronts of DIPs and IPs allows DIPs to infect a greater number of target cells, while the front is constantly led by IPs.

### DIPs efficiency depends on maturation rate of co-infected cells

Increase in the maturation rate of co-infected cells in *E*_D_ class increases the travelling speed of DIPs so that DIPs compete with IPs at the front. In case of equal maturation rates (*γ* = *γ*_D_), the travelling front of DIPs go along IPs by co-infecting cells of *E* class, see Figure 2. As gamma is increased, and the fraction of DIPs produced by cells in *I*_D_ class is chosen sufficiently high, the difference in the travelling fronts of DIPs and IPs is reduced, see Figure 6(c). Consequently, DIPs start to infect target cells and therefore the peak value of IPs, that is lying near the front, sufficiently decreases as shown in Figure 6(a) and (b). However, since the travelling front is dominated by IPs, the speed of infection spread remains same as in the above results. This provides an interesting comparison to a recent study in which IPs spread is substantially reduced with the presence of DIPs. Thus, the close proximity between the DIPs and IPs requires stochastic effects at the travelling front in order to bring delays in the virus spread (Baltes et al., 2017) which brings a challenge to be addressed by deterministic modeling.

**Figure 6:**
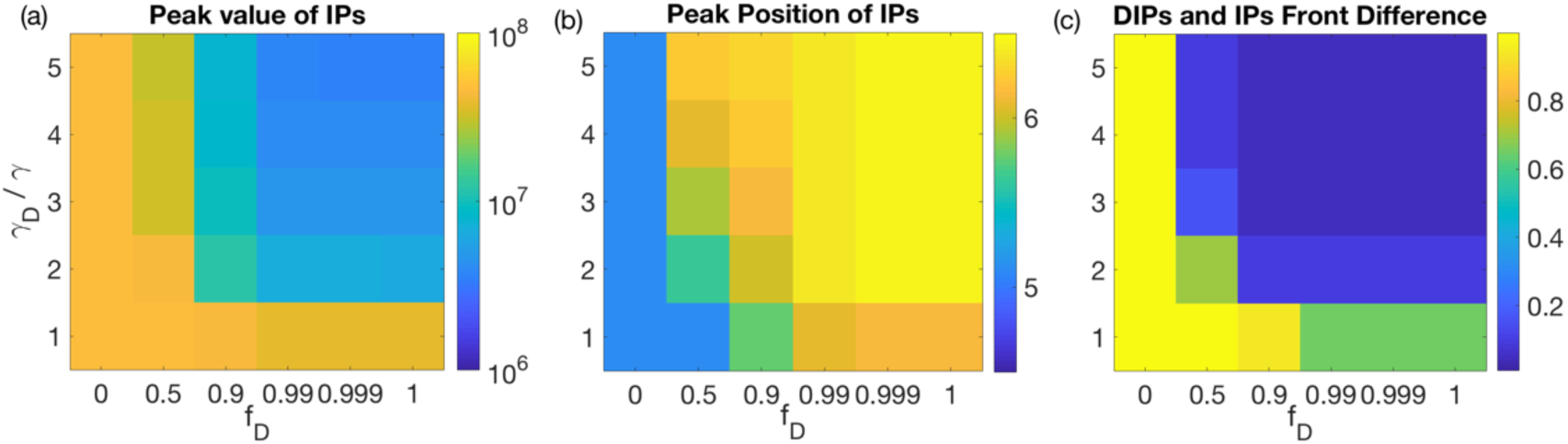
Heatmap for the parameter space, fraction increase in the maturate rate of the cells in *E*_D_ class (m = *γ*_D_ / *γ)* and fraction of DIPs produced by the cells in *I*_D_ class (*f*_D_), at final time t = 25 hr. Subplot (a) explains peak value of IPs, (b) describes the difference between the front positions of DIPs and IPs and (c) shows the spatial position of IPs at the peak value.

## Discussion

This study describes a complex dynamical system of with-in host virus infection that involves time-dependent reaction processes, incorporating spatial distributions of cells and virus particles. We explain the role of DIPs in reducing the number of virus particles when a virus infection spreads and proceeds within the tissues where the cells are arranged in a group or a layer (Murillo et al., 2013). In the conventional antiviral drug therapies, non-spatial dynamics of homogeneously mixed populations were designed to directly control the infections by reacting with key viral components. Defective interfering particles (DIP) have shown a distinctive behavior as compared to the classical antiviral agents. These particles have shown a substantial decrease in the virus growth without any direct interaction to the wild-type viruses, i.e. IPs, (Baltes et al., 2017).

The parasitic behavior of DIPs reveals its quality of natural spread along the IPs. Our results show that the transformation of naturally occurring defective particles by sophisticated engineering techniques can make them propagate along IPs on the spatially distributed target cells to continuously interfere in the infection process as shown in Figure 2(b). However, its existence is contingent to its production efficacy and its presence along the wild-type viruses on the locally distributed target cells, see for example Figure 2(a) and (c). Therefore, it is important for the existence of DIP to propagate along IPs at the same speed as the speed of spread of wild-type virus.

The transient phase of viral infection is crucial for DIPs to play their role in suppressing the infection at the within host level. The only objective of DIPs is to infect maximum number of non-infected target cells rather than co-infecting those cells that are in the eclipse phase of IP infection. This is necessary for the effective survival of DIPs when the travelling front of IPs is established. We assert that increase in the DIPs infection to non-infected target cells is contingent to the improvement of multiple factors such as fraction of DIPs production (*f*_D_), virions production rate (*p*_D_), cells maturation rate (γ_D_) and the initial conditions (ICs) for virus particles, i.e. maximum initial concentration (*V*_D0_) and ratio between the spread of DIPs and IPs (*σ*_D_ / *σ*). Our results show that when the fraction of DIPs production (*f*_D_) from co-infected cells is high, DIPs are able to infect only those cells that are in the eclipse phase of IP infection (see for example Figure 2(b)). In this way, the spread of both particles, IPs and DIPs, depends on the IP infected cells rather than the target cells. It may create an unfavorable situation for DIPs so that they are unable to even lag behind IPs. Although a very high initial condition for DIP is able to suppress the wild-type infection process (see Figure 2(c) and Figure 3(c)) but a biologically reasonable initial condition indicate that DIP are unable to infect target cells during the steady behavior of the travelling front of IPs (see Figure 3(g) and (h)). Moreover, there is a visible decrease in the concentration of IPs (see Figure 4(a)) while the peak virus concentration shifts near the front position (Figure 4(b)). It is interesting, but not surprising, that no effect has been observed on the speed of the traveling front of IPs (see Figure 3(a) and (b) and Figure 4(c)). To further enhance the role of DIPs in suppressing the IPs, other factors are necessary to be involved.

The increase in the production rate and decrease in the maturation rate of co-infected cells increases speed of the travelling front of DIPs. Consequently, the travelling fronts of IPs and DIPs come closer to each other. We noted that the existence of efficient DIPs in the infected region with same travelling speed as IPs delayed IPs to reach their maximum peak concentration (see Figure 3(a) and (b)) and brought their peak position near the travelling front (see Figure 3(d) and (e)). Interestingly, the speed of IPs spread at the front remained constant even when the front difference of DIPs and IPs was very small (see Figure 3(g), (h) and Figure 5(c), (d)). This provides a good contrast with a previous study which has demonstrated a considerable delay in the spread of infectious particles (Baltes et al., 2017), suggesting that the delays are due to the occurrence of stochastic events at the travelling front when the DIPs are very close to IPs.

Current engineering designs of DIP introduced it as a dual-purpose strategy that can play an important role in reducing viral load like a prodrug as well as generating strong immune signals at the infected region like a pro-vaccine (Baltes et al., 2017; Dimmock and Easton, 2014; Tapia et al., 2013). This study focused on the prodrug features of DIP that reduced the concentration of IPs within-host while giving a hope that its pro-vaccine features can be addressed by including its role in inducing IFN signals (Panda et al., 2010; Tapia et al., 2013).

Recent advancements have introduced a new class of segmented viruses that has incomplete set of genome segments, called as semi-infectious particles (SIPs). These particles are non-infectious under traditional limiting-dilution assays (Brooke et al., 2013) while they become infectious through multiplicity reactivation, when MOI is high (Brooke, 2014). Therefore, it would be interesting to see how SIPs are able to traverse the front position of wild-type viruses and able to bring DIPs close to their travelling front. The stochastic effects to the DIP spread on the target cells near the travelling front may show some interesting outcomes in terms of reducing the speed of virus spread.

## Acknowledgement

I am thankful to Alex Farrell for his discussions and feedback about this work. I am also thankful to Michael Gary Lavigne for his help regarding linguistics and technical words. This work was funded by DARPA, USA.

## Contributions

This report is solely written by Qasim Ali, not revised by Ruian Ke. The results are discussed with Ruian Ke mostly by email exchange.

